# pyseer: a comprehensive tool for microbial pangenome-wide association studies

**DOI:** 10.1101/266312

**Authors:** John A Lees, Marco Galardini, Stephen D Bentley, Jeffrey N Weiser, Jukka Corander

**Affiliations:** Department of Microbiology, New York University School of Medicine, New York, NY, 10016 USA; European Bioinformatics Institute (EMBL-EBI), Hinxton, CB10 1SD, United Kingdom; Infection Genomics, Wellcome Sanger Institute, Hinxton, CB10 1SA, United Kingdom; Department of Biostatistics, University of Oslo, 0372 Oslo, Norway; Helsinki Institute of Information Technology (HIIT), Department of Mathematics and Statistics, University of Helsinki, 00014 Helsinki, Finland

## Abstract

**Summary:** Genome-wide association studies (GWAS) in microbes face different challenges to eukaryotes and have been addressed by a number of different methods. *pyseer* brings these techniques together in one package tailored to microbial GWAS, allows greater flexibility of the input data used, and adds new methods to interpret the association results.

**Availability and Implementation:** *pyseer* is written in python and is freely available at https://github.com/mgalardini/pyseer, or can be installed through *pip*. Documentation and a tutorial are available at http://pyseer.readthedocs.io.

**Supplementary information:** Supplementary data are available online.

## 1 Introduction

Finding genetic variation associated with bacterial phenotypes such as antibiotic resistance, virulence and host specificity has great potential for a better understanding of the evolution of these traits, and may be able to inform new clinical interventions. Genome-wide association studies (GWAS) address this question in a hypothesis-free manner, and also provide a framework in which to quantify the amount of phenotypic variation due to the genetics of the bacterium. The recent availability of thousands of whole-genome sequences from bacterial populations has made this approach possible, though issues of strong clonal population structure and a variable pangenome must be accounted for in any successful analysis (Power *et al*., 2016).

One method that addresses these issues in a scalable manner is *SEER* (Lees *et al*., 2016), which uses variable length k-mers as a generalized variant to represent variation across the pangenome and linear models with a control for population structure to perform the association. Other methods include: *bugwas*, which uses a linear mixed model (LMM) and also checks for lineage effects (Earle *et al*., 2016); *scoary*, which tests clusters of orthologous genes (COGs) with and without accounting for population structure (Brynildsrud *et al*., 2016); and phylogenetic regression, which uses a known phylogeny to adjust for the covariance between samples (Garland and Ives, 2000).

Each of these methods has its own set of advantages and limitations, so recent GWAS analyses have used a combination of these techniques along with methods designed for human genetics, often in a somewhat ad-hoc manner that requires tedious file format conversion and familiarity with a wide range of methods and software packages (Desjardins *et al*., 2016; Lees *et al*., 2017).

As the number of large bacterial population datasets increases, we recognize the need to make these methods more accessible to the field. We have therefore re-implemented *SEER* in an easy-to-use and install *python* package, *pyseer*. *pyseer* also includes many additional features covering new association models, input sources and output processing. This brings together all the methods mentioned above into a single piece of software, and further enables new combinations of analysis. Our method is equally applicable to viral GWAS, and indeed any species with structured populations and/or genomes with significant variation in gene content.

## 2 Methods

The foundation of *pyseer* is a direct python re-implementation of *SEER* (originally written in C++). K-mers of variable length counted from draft assemblies are used as the input, and their association with a phenotype of interest is assessed by fitting a generalized linear model to each k-mer. To control for population structure, multi-dimensional scaling of a pairwise distance matrix is performed and these components are included as fixed effects in each regression. In the case of a binary phenotype with a highly penetrant variant (those with high effect sizes), Firth regression (Heinze and Schemper, 2002) is performed to maintain power. Significant k-mers, after adjusting for multiple testing, can then be mapped to a reference annotation to produce a Manhattan plot and find regions of the genome associated with the phenotype.

After re-implementation, results were the same as *SEER* (Supplementary figure 1). *pyseer* is of comparable speed, with the exception of a continuous phenotype with fixed effects (Table 1). We then expanded the features in *pyseer* to bring together the different methods mentioned above.

**Table 1.**
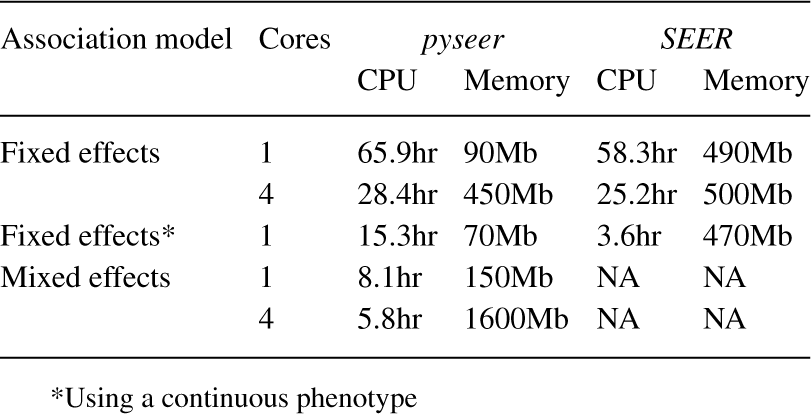
Resource comparison between pyseer and SEER using the tutorial dataset (15.1M k-mers, no filtering).

### 2.1 Input sources

In addition to k-mers, the original focus of *SEER*, it can be convenient to test for association of SNPs and INDELs called against a reference genome. These can be used to confirm the k-mer results, as a first pass to confirm the analysis chosen is working correctly or to test for association of close or adjacent SNPs, which would be split into many low frequency k-mers. Presence or absence of COGs and aligned intergenic regions can also be useful, as shown by *scoary* (Brynildsrud *et al*., 2016). The consequence of these variants can be predicted (e.g. synonymous, frameshifting) which requires more downstream processing for k-mers.

*pyseer* can natively read all these types of variant from VCF or Rtab files, which also allows for any user defined input type (for example copy number variants). We also enable grouping of variants by genomic region to perform burden testing. This now allows users to perform analysis of rare variation while still accounting for population structure.

An important technical difficulty of using *SEER*, and most common linear mixed model implementations, is making sure that the membership and order of samples between the variant calls, phenotype file and population structure all match up correctly. Many of the methods do not check for this, and can run an incorrect analysis without producing a warning. In *pyseer* this label matching is done automatically, and the intersection of samples used is reported to the user.

### 2.2 Association models

*pyseer* implements the same fixed effect generalized linear regression model as *SEER*, including Firth regression. We have added two new multidimensional scaling algorithms in *pyseer*, and a more streamlined interface with mash (Ondov *et al*., 2016) to compute population structure. Alternatively, when a high quality phylogeny is available, *pyseer* can use this instead to adjust for population structure in a manner analogous to phylogenetic regression.

As a major alternative we have included an LMM, which uses the kinship matrix to estimate random effects. This model has been shown to control population structure across a range of scenarios. We have used the FaST-LMM implementation, which allows association in linear time (Lippert *et al*., 2011), along with a kinship matrix estimated from a subset of variants or from a phylogeny. This allows association of all k-mers under the mixed model in a few hours, which was previously computationally and bioinformatically challenging.

We have also included a method to estimate possible lineage effects, based on the procedure used in *bugwas*. After association with the phenotype, variants are also associated with lineages (which can be determined by *pyseer* or defined by the user). Those variants which are associated with both the phenotype and with a lineage associated with the phenotype can be prioritised for further analysis outside of GWAS.

### 2.3 Output processing

It has been suggested that the number of unique variant patterns is a sensible way to set the multiple testing threshold (Earle *et al*., 2016), however these can be difficult to count due to the size of k-mer data. *pyseer* uses sorting of hashes to efficiently calculate this threshold.

Finally, we have added tools to make interpretation of significant kmers more streamlined. The user can firstly map their results against any number of reference genomes for interactive display in *phandango* (Hadfield *et al*., 2017) (Supplementary figure 2). We have also added the ability to summarise k-mer results at the gene level through iteratively mapping to reference and draft annotations, which can be used to show complementary information about effect size, coverage and minor allele frequency (Supplementary figure 3).

## 3 Discussion

Starting with a re-implementation of *SEER*, we have added the models and input types used by other microbial and human GWAS approaches into a single package. Analyses which were previously challenging to perform, such as association of all variable length k-mers with a LMM, can be performed in an efficient and user-friendly manner. We have also enabled new types of analysis, such as population structure corrected burden testing, and gene level summaries of kmers. Our package includes comprehensive documentation and a tutorial (http://pyseer.readthedocs.io/en) which shows how to perform GWAS using the new input sources and both association models, as well as how to interpret significant k-mers. We have implemented unit tests in our code to ensure consistency of output as features are added.

With *pyseer* we have therefore reconciled many of the existing methods for regression-based microbial GWAS into a single package. Our focus on documentation and ease-of-use of *pyseer* will make GWAS more accessible to the microbial genomics community.

## Acknowledgements

We wish to thank Leonor Sanchez-Buso for an early version of the k-mer annotation script. We would also like to thank Pedro Beltrao and Chrispin Chaguza for constructive comments.

